# Reply: Evidence that *APP* gene copy number changes reflect recombinant vector contamination

**DOI:** 10.1101/730291

**Authors:** Ming-Hsiang Lee, Christine S. Liu, Yunjiao Zhu, Gwendolyn E. Kaeser, Richard Rivera, William J. Romanow, Yasuyuki Kihara, Jerold Chun

## Abstract

In the accompanying comment^1^, Kim *et al*. concluded that somatic gene recombination (**SGR**) and amyloid precursor protein (***APP***) genomic complementary DNAs (**gencDNAs**) in brain are contamination artifacts and do not naturally exist. We disagree. Here we address the three types of analyses used by *Kim et al*. to reach their conclusions: informatic contaminant identification, plasmid PCR, and single-cell sequencing. Additionally, Kim *et al*. requested “reads supporting novel *APP* insertion breakpoints,” and we now provide 10 different examples that support *APP* gencDNA insertion within eight chromosomes beyond wildtype *APP* on chromosome 21 from Alzheimer’s disease (**AD**) samples. If SGR exists as experimentally supported here and previously^2,3^, contamination scenarios become moot. Our informatic analyses of data generated by an independent laboratory (Park *et al.)^4^*, complement and are entirely consistent with what Lee *et al*.^2^ presented via nine distinct lines of evidence, in addition to three from a prior publication^3^. Plasmid contamination was identified in a single pull-down dataset after publication of Lee *et al*.^2^; however subsequent analyses did not alter any of our conclusions including those of our prior publications^3,5^ and plasmid contamination-free replication of this approach by ourselves and others supported the original conclusions. Novel retro-insertion sites, alterations of *APP* gencDNA number and form with cell type from the same brain and pathogenic SNVs occurring only in AD, all support the existence of *APP* gencDNAs produced by SGR.

## Identification of novel *APP* gencDNA insertion sites

One predicted outcome of SGR is the generation of novel retro-insertion sites distinct from the wildtype locus, as we demonstrated using DNA *in situ* hybridization (**DISH**; Lee *et al*. Figure 2n). Analyses of independently published datasets (Park *et al*.)^4^ produced by whole-exome pull-down of DNA from laser-captured hippocampus or blood revealed 10 different *APP* insertion sites within eight different chromosomes (**Figure 1, Supplementary Table 1**). We identified clipped reads spanning *APP* UTRs and novel genomic insertion sites on chromosomes 1, 3, 9, 10, and 12 (**Figure 1a**; wildtype *APP* is located on chromosome 21). The corresponding paired-end reads mapped to the same inserted chromosome. We also identified reads spanning *APP* exon::exon junctions of gencDNAs that had mate-reads mapping to other genomic sites on chromosomes 1, 3, 5, 6, and 13 (**Figure 1b**). We are unaware of contamination sources capable of producing these results that are entirely consistent with our DISH data showing *APP* gencDNA locations distinct from wildtype *APP*. These novel *APP* gencDNA insertion sites strongly support the natural occurrence of *APP* gencDNAs.

**Figure 1.**
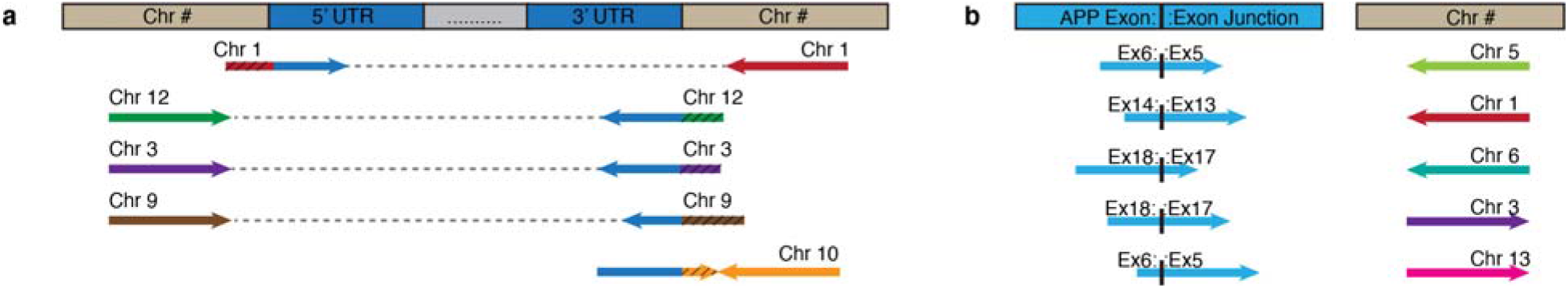
Identification of novel APP insertion sites in the human genome. a) Clipped reads spanning *APP* UTRs and novel chromosomal insertion sites were identified. The paired matereads of the clipped reads (black stripes) uniquely mapped to the same chromosomes. b) Discordant read-pairs were identified where one read spanned an *APP* exon::exon junction and the corresponding mate-read mapped to a novel chromosome. Each chromosome has a unique color. Arrowhead direction represents the read orientation after mapping to the human reference genome. Arrows oriented in the same direction support sequence inversions. See detailed sequence and alignment information in Supplementary Table 1.

**Figure 2.**
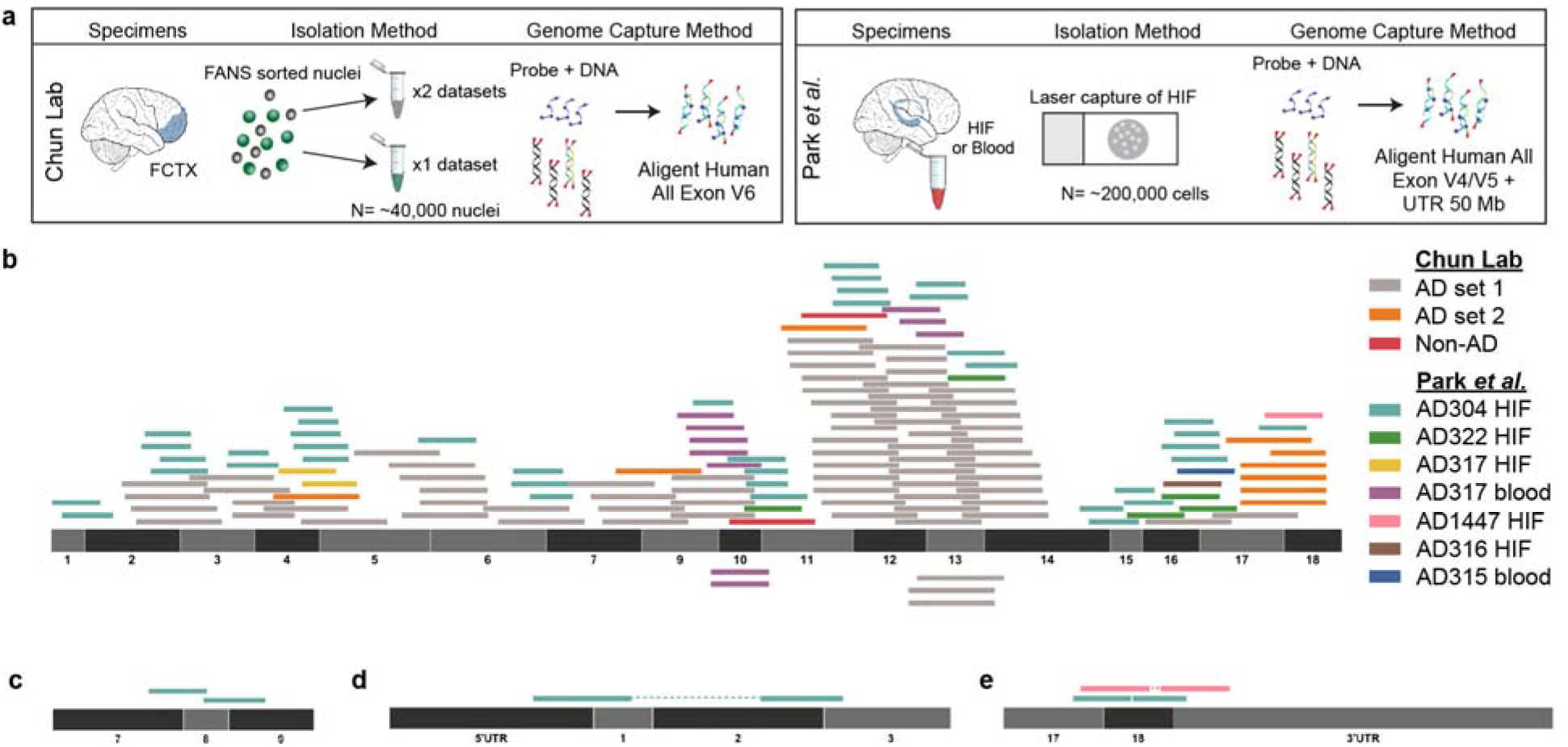
Identification of *APP* gencDNA sequences in 10 new whole-exome pull-down datasets from two independent laboratories. a) Method schematic depicting the standard protocol for whole-exome pull-downs and highlighted methodological differences between the independent laboratories are presented. b) *APP-751* sequence with non-duplicate gencDNA reads from the 10 new datasets; color key indicates the source reads for all panels. c) Reads mapping to junctions between *APP* exons 7, 8, and 9 that are absent from *APP-751*. d,e) Paired reads that represent a DNA fragment containing both an exon::exon junction and an *APP* 3’ or 5’ UTR.

An *APP* plasmid contaminant (pGEM-T Easy *APP*) was found in our single pull-down dataset, however we could not definitively determine which *APP* exon::exon reads were due to gencDNAs vs. plasmid contamination, especially in view of the 11 other distinct and uncontaminated approaches that had independently supported and/or identified *APP* gencDNAs. Three other pull-down datasets from our laboratory were informatically analyzed and found to contain *APP* gencDNA reads while being free from *APP* plasmid contamination by both VecScreen^6^ and subsequent use of Kim *et al.’s* Vecuum script^7^ (**Figure 2a,b**). Possible external source contamination noted by Kim *et al*. in two of three datasets could not definitively account for all *APP* exon::exon junctions.

The recent availability of independently generated AD datasets^4^ provided a test for the reproducibility of *APP* gencDNA identification. Five different sporadic AD (**SAD**) brains and two AD blood samples contained *APP* gencDNA sequences and were plasmid-free by Vecuum^7^ screening (**Figure 2a-e**). In addition to exon::exon junction reads and novel insertion sites, we also identified *APP* UTR sequences paired with reads containing *APP* gencDNA exon::exon junctions (**Figure 2d,e**). This may be explained by a key experimental design factor: Park *et al*.’s pull-down probes contain sequences corresponding to *APP* 5’ and 3’ UTRs.

In addition to *APP* plasmid and amplicon contaminants, Kim *et al*. invoked genome-wide mouse and human mRNA contamination in the Park *et al*. dataset. We cannot address conditions in the Park *et al*. laboratory but note that it is completely independent of our own. Kim *et al*.’s explanation implicates the generation of DNA from mRNA: a process that requires reverse transcriptase activity. The Agilent SureSelect pull-down employed by Park *et al*. and in our experiments do not use reverse transcriptase (**Figure 2a and Supplementary Methods**), and we are unaware of any mechanism that would generate DNA from RNA in the absence of reverse transcriptase activity under the employed conditions. An alternative explanation is the existence of gencDNAs affecting other genes as we previously detected in non-*APP* intra-exonic junctions (**IEJs**) found in commercial cDNA Iso-Seq datasets (**Extended Data Figure 1**). Additional validation would be required for new genes, however we note that an average of 450 megabase-pairs of extra DNA exist within AD neurons^3^ that could accommodate new gencDNA sequences. Kim *et al*. further invoked genome-wide mouse and human mRNA contamination in the Park *et al*. dataset to account for *APP* gencDNAs, an explanation conflicting with available data. Mouse-specific single nucleotide polymorphisms (**SNPs**) in the Park *et al*. dataset cannot account for all *APP* gencDNA-supporting reads: five of seven *APP* exon::exon junction sequences do not contain putative mouse-specific SNPs at the specific region reported by Kim *et al*. (**Figure 3**; Kim *et al*. Figure 2d). Most critically, novel *APP* gencDNA insertion sites identified here cannot be explained by genome-wide mRNA contamination.

**Figure 3.**
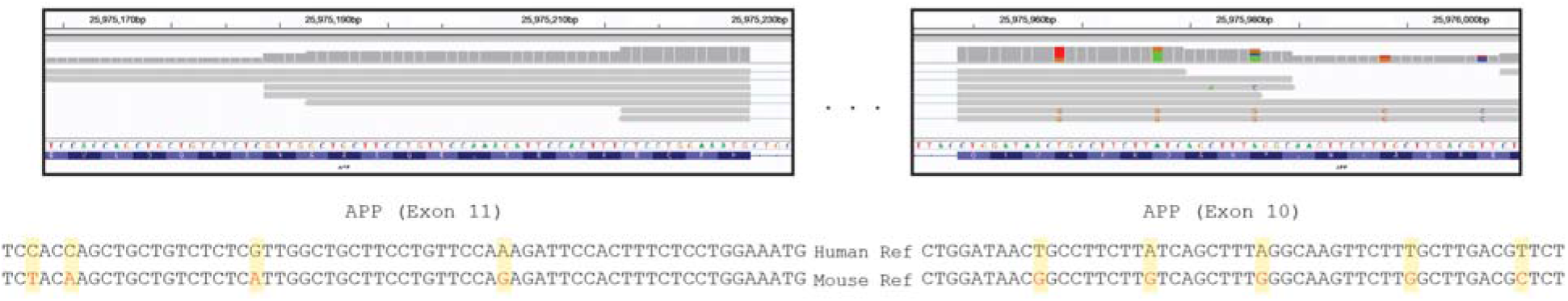
Five *APP* gencDNA-supporting reads spanning exon::exon junctions that do not contain mouse-specific SNPs. *APP* gencDNA reads were identified that span the *APP* exon10::exon11 junction from the Park *et al*. datasets. The reference sequences of human and mouse exons are indicated and the positions where the nucleotides differ are highlighted. Five of the seven exon::exon junction-spanning reads do not contain mouse-specific SNPs.

## Non-biological data are generated by PCR of *APP* plasmids in Kim *et al*

Kim *et al*. used PCR of *APP* splice variant plasmids which generated sequences containing IEJs. However, multiple discrepancies in this approach and results differ from our biological IEJs and gencDNAs: 1) experimental conditions beyond our primer sequences were different: Kim *et al*. employed twice the concentration of primers and >1 million times more template (250 picograms of *APP* plasmid is 4.6 x 10^7^ copies vs. ~40 gencDNA copies in our PCR of 20 nuclei (based on Lee *et al*. Figure 5^2^: DISH 16/17 averaged ~1.8 copies/SAD nucleus)); 2) both gencDNA and IEJ sequences can be detected with as few as 30 cycles of PCR as we used in single molecule real-time (**SMRT**) sequencing (Lee *et al*. Figure 3)^2^ vs. 40 cycles used by Kim *et al*.; 3) agarose gels in Kim *et al*. are uniformly and unambiguously dominated by a vastly over-amplified ~2 kb band (Kim *et al*. Figure 1c and Extended Data Figure 3a) that is never seen in human neurons despite our routine identification of myriad smaller bands (*c.f*., Lee *et al*. Figure 2b)^2^. We did observe an over-amplified ~2 kb band in our purposeful plasmid transfection experiments that also utilized PCR; however, gencDNA and IEJ formation was comparatively limited, and critically, required both reverse transcriptase activity and DNA strand breakage (Lee *et al*., Figure 4^2^); and 4) only 45 unique IEJs from AD and 20 from non-diseased brains were identified (Lee *et al*. Figure 3 with some overlap, fewer than 65 total)^2^ compared to the 12,426 identified by Kim *et al*. (~200-fold increase over biological IEJs; Kim *et al*. Supplementary Table 1). We wish to note that microhomology regions within *APP* exons are intrinsic to *APP’s* DNA sequence and that microhomology mediated repair mechanisms involve DNA polymerases^8,9^. Kim *et al.’s* PCR results differ from our biological data yet may inadvertently support endogenous formation of at least some IEJs within DNA rather than requiring RNA.

## Detection of IEJs without use of *APP* PCR

Despite these differences between the non-biological plasmid PCR data generated by Kim *et al*. and our data, Kim *et al*. concludes that IEJs from our original study^2^ might have originated from contaminants. To eliminate this possibility, Lee *et al*.^2^ presented four lines of evidence for *APP* gencDNAs containing IEJs that are independent of *APP* PCR: two different commercially produced cDNA SMRT sequencing libraries, DISH, and RNA *in situ* hybridization (**RISH**). The SMRT sequencing libraries revealed IEJs within *APP* (Lee *et al*. Extended Data Figure 1E)^2^ as well as other genes (**Extended Data Figure 1**), which cannot be attributed to plasmid contamination or PCR amplification. DISH and RISH results support the existence of *APP* gencDNAs and IEJs (see **Supplementary Discussion** and Lee *et al*., Figure 2, Extended Data Figures 1 and 2)^2^ by using custom-designed and validated commercial probe technology (Advanced Cell Diagnostics, ACD), which was independently shown to detect exon::exon junctions^10^ and single nucleotide mutations^11^. Thus, gencDNAs and IEJs are detectable in the absence of targeted PCR. Importantly, the contamination proposed by Kim *et al*. cannot account for the dramatic change in the number and forms of *APP* gencDNAs occurring with disease state. The change is also apparent when comparing cell types, where signals are vastly more prevalent in SAD neurons compared to non-neurons from the same brain and processed at the same time by DISH (Lee *et al*. Figures 5)^2^. Independent PNA-FISH and dual-point-paint experiments from our previous work further support *APP* gencDNAs^3^ (**Table 1**). Critically, SMRT sequencing identified 11 single nucleotide variations that are considered pathogenic in familial AD, which were only present in our SAD samples, none of which exist as plasmids in our laboratory.

**Table 1.** Summary of targeted and non-targeted *APP* PCR methods and lines of evidence that support *APP* gencDNAs and IEJs.

|  | Method | Targeted *APP* PCR | Support for the existence of IEJs and gencDNAs | Reference |
| --- | --- | --- | --- | --- |
| **Approaches without targeted *APP* PCR** |  |  |  |  |
| 1 | RISH on IEJ 3/16 | None | IEJ 3/16 RNA signal is present in human SAD brain tissue | Lee *et al.* |
| 2 | Whole Transcriptome SMRT sequencing | None | An independent commercial source identified IEJs in *APP* and other genes | Public data<sup>1</sup>, Lee *et al.*, This Reply |
| 3 | Targeted RNA SMRT sequencing | None | RNA pull-down that identified *APP* IEJs | Public data set<sup>1</sup>, Lee *et al.* |
| 4 | DISH of gencDNAs | None | IEJ 3/16 and exon::exon junction 16/17 showed increases in SAD neurons compared to non-neurons from the same brain and non-diseased neurons; J20 mice containing the *APP* transgene under a PDGF-β-promoter show increased number and size of signal compared to non-neurons and WT mice | Lee *et al.* |
| 5 | Dual point-paint FISH | None | Identified *APP* CNVs of variable puncta size that were not always associated with Chr21 | Bushman *et al.* |
| 6 | PNA-FISH | None | *APP* exon copy number increases show variable signal size and shape with semi-quantitative exonic probes | Bushman *et al.* |
| 7 | Agilent SureSelect Targeted pull-down | None | Identified *APP* gencDNAs in SAD brains; contains plasmid sequence contamination | Lee *et al.*, This Reply |
| New #7 | Agilent All-Exon pull-down | None | All-Exon pull-downs with no plasmid contamination by Vecuum contain *APP* gencDNA sequences and evidence of gencDNA UTRs and novel insertion sites | Park *et al.*, This Reply |
| **Approaches with targeted *APP* PCR** |  |  |  |  |
| 8 | RT-PCR and Sanger sequencing | Oligo-dT primed and targeted *APP* primers | Novel *APP* RNA variants with IEJs; predominantly in neurons from SAD brains | Lee *et al.* |
| 9 | Genomic DNA PCR and Sanger Sequencing | Yes | Identified *APP* gencDNAs with IEJs; predominantly in neurons from SAD brains | Lee *et al.* |
| 10 | Genomic DNA PCR and SMRT sequencing | Yes | IEJ/gencDNAs were more prevalent in number and form in SAD neurons compared to non-diseased neurons; Identified 11 pathogenic SNVs that were only present in SAD samples | Lee *et al.* |
| 11 | *APP*-751 over-expression in CHO cells | Yes | IEJ and gencDNA formation required DNA strand breakage and reverse transcriptase | Lee *et al.* |
| 12 | Single-cell qPCR | Yes; individual exon | Intragenic exon 14 single-cell qPCR showed copy number increases in SAD prefrontal cortical neurons over cerebellar neurons from the same brain | Bushman *et al.* |
| CNV, copy number variation; DISH, DNA *in situ* hybridization; FISH, fluorescence *in situ* hybridization; IEJ, intra-exonic junction; PNA, peptide nucleic acid; RISH, RNA *in situ* hybridization; SAD, sporadic Alzheimer's disease; SMRT, single molecule real-time. |  |  |  |  |
| <sup>1</sup>The Alzheimer brain Iso-Seq dataset was generated by Pacific Biosciences, Menlo Park, California. Additional sequencing information and analysis is provided at https://downloads.pacbcloud.com/public/dataset/Alzheimer_IsoSeq_2016/. |  |  |  |  |

Kim *et al*. compared *APP* gencDNA copy number estimates from pull-down sequencing and DISH. However, a direct comparison is not possible since the two methodologies are fundamentally different. For example, pull-downs employ solution hybridization on isolated DNA, while DISH uses solid-phase hybridization on fixed and sorted single nuclei. Moreover, the sequences targeted between the two are not the same. Pull-down probes target wildtype sequences for endogenous and gencDNA loci, resulting in pull-down competition. By contrast, DISH probes target only gencDNA sequences to provide greater sensitivity. Competition by wildtype loci reduced the efficiency of capture, which is underscored by 32% to 40% of nuclei that do not contain gencDNAs and would contribute only wildtype sequences (Lee *et al*., Figure 5c,f). Moreover, a majority of gencDNA positive nuclei (62% to 73%) showed two or fewer signals (Lee *et al*., Figure 5c,f) which reduced the relative representation of gencDNA loci. Since IEJs do not contain the full exon sequence, there is inefficient hybridization and a lack of sequence capture and detection. This limitation is overcome by SMRT sequencing (**Extended Data Figure 1** and Lee *et al*., Extended Data Figure 1e). Lastly, multiple other protocol variations exist which explain the hypothesized discrepancies including tissue preparation, fixation, and hybridization conditions.

## Single-cell whole-genome sequencing limitations may prevent *APP* gencDNA detection

Kim *et al.’s* third type of analysis yielded a negative result via interrogation of their own singlecell whole-genome sequencing (**scWGS**) data, which cannot disprove the existence of *APP* gencDNAs. An average of nine neurons from seven SAD brains were examined, raising immediate sampling issues required to detect mosaic *APP* gencDNAs. Kim *et al*. identified “uneven genome amplification” (Kim *et al*. and ^12–14^) producing ~20% of the single-cell genome having less than 10X depth of coverage^14^ with potential amplification failure at one (~9% allelic dropout rate) or both alleles (~2.3% locus dropout rate)^12,14^. These limitations are compounded by potential amplification biases reflected by whole-genome amplification failure rates that may miss neuronal subtypes and/or disease states, which is especially relevant to single copies of *APP* gencDNAs that are as small as ~0.15 kb (but still detectable by DISH). Kim *et al*. state that the increased exonic read depth relative to introns reliably detects germline retrogene insertions in single cells from affected individuals (Kim *et al*., Figure 3b); however, these data also demonstrate that increased exonic read depth is *not* observed in all cells – or even a majority in some cases – from the same individuals carrying the germline insertions of SKA3 (AD3 and AD4) and ZNF100 (AD2). These results demonstrate inherent technical limitations in Kim *et al*. that prevent accurate detection of even germline pseudogenes present in all cells, thus explaining an inability to detect the rarer mosaic gencDNAs produced by SGR. Kim *et al.’s* informatic analysis is also based on the unproven assumption that gencDNA structural features are shared with processed pseudogenes and LINE1 elements (Kim *et al*. Figure 3a and Extended Data Figure 1a), and possible differences could prevent straightforward detection under even ideal conditions as has been documented for LINE1^15^. These issues could explain Kim *et al.’s* negative results.

Considering these points, we believe that our data and conclusions supporting SGR and *APP* gencDNAs remain intact and warrant their continued study in the normal and diseased brain.

## Author Contributions

MHL, YK, and WR conducted laboratory experiments; CSL and YZ analyzed sequencing data; and JC conceived and oversaw the experiments. All authors wrote and edited the manuscript. This <u>Reply</u> was the work of current laboratory members.

## Data availability

Data from Park *et al*. were deposited in the National Center for Biotechnology Information Sequence Read Archive database, accession number PRJNA532465. Data from the newly reported full exome pull-down datasets will be provided for the *APP* locus upon request.

## Code availability

The source codes of the customized algorithms are available on GitHub https://github.com/christine-liu/exonjunction.

## Acknowledgements

We thank Dr. Laura Wolszon and Ms. Danielle Jones for manuscript editing. Research reported in this publication was supported by the NIA of the National Institutes of Health under award number R56AG067489 and P50AG005131 (J.C.); and NINDS R01NS103940 (Y.K.). This work was supported by non-Federal funds from The Shaffer Family Foundation, The Bruce Ford & Anne Smith Bundy Foundation, and Sanford Burnham Prebys Medical Discovery Institute funds (J.C.). The content is solely the responsibility of the authors and does not necessarily represent the official views of the National Institutes of Health.

---

**Extended Data Figure 1.**
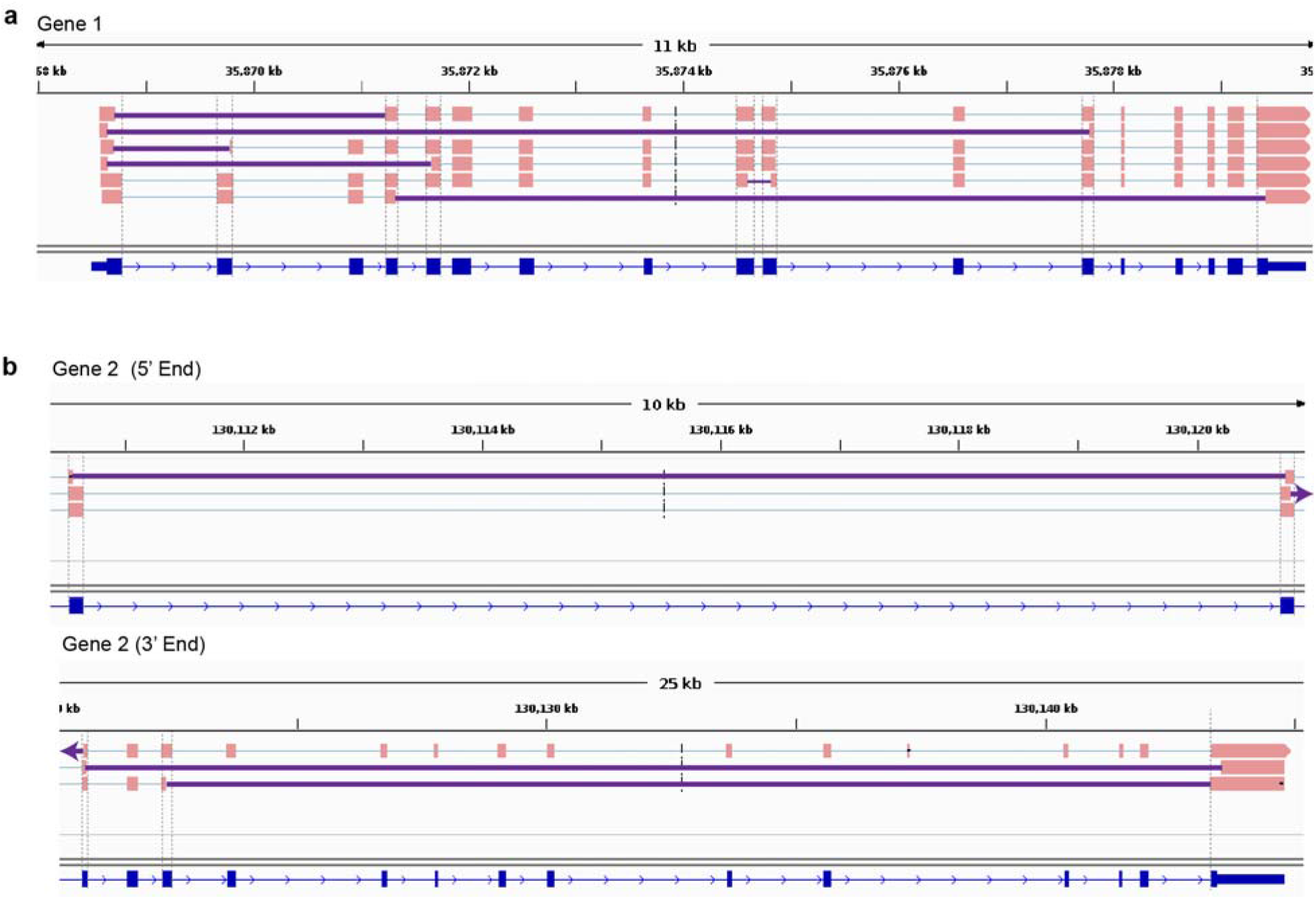
IEJs identified from commercially available long-read transcriptome datasets in two genes other than *APP*. Sequences containing IEJs were identified and shown for a) Gene 1 and b) Gene 2. Gene 2 is shown in two parts. Grey dashed lines show ends of RefSeq exons; solid purple lines denote IEJs. All splice isoforms were examined. The Alzheimer brain Iso-Seq dataset was generated by Pacific Biosciences, Menlo Park, California, and additional information about the sequencing and analysis is provided at https://downloads.pacbcloud.com/public/dataset/Alzheimer_IsoSeq_2016/.

