## Supplementary Information for "Reply: Evidence that *APP* gene copy number changes reflect recombinant vector contamination"

**Supplementary Methods**

**Agilent Human All Exon V6 pull-down experimental protocol and analysis**

**Human brain tissue**

IRB-approved fresh frozen human brain tissue was provided by the University of California San Diego (UCSD) Alzheimer’s Disease Research Center (ADRC) and the University of California Irvine (UCI) Institute for Mind Impairments and Neurological Disorders (MIND).

**Nuclei extraction and fluorescence-activated nuclear sorting (FANS)**

Human brain nuclei were isolated as described previously^1^. Prior to human brain nuclei sorting, nuclei were labeled with anti-NeuN rabbit monoclonal antibody (1:800) (Millipore, Germany) and Alexa Fluor 488 donkey anti-rabbit IgG (1:500) (Life Technologies, Carlsbad, CA), and counterstained with propidium iodide (50μg/ml) (Sigma, St. Louis, MO). RNAse A (100ug/ml) was included in all subsequent steps after initial nuclei isolation, including primary and secondary antibody incubations. Diploid NeuN-positive and negative nuclei were gated by PI and Alexa Fluor 488 fluorescence and sorted into appropriate populations for all human exon pull-down experiments. FANS was performed on a FACSAria Fusion (BD Biosciences, Franklin Lakes, NJ).

**DNA extraction**

DNA was extracted from isolated neuronal nuclei populations and precipitated with isopropanol. Briefly, nuclei were incubated with proteinase K in 550 μl PK buffer (50mM Tris pH8.0, 0.1M EDTA, 0.1M NaCl, 1% SDS) overnight at 55°C. Samples were then treated with an RNase cocktail enzyme mix (ThermoFisher, Waltham, MA) for 2 hr, followed by the addition of 250 μl saturated NaCl. After centrifugation, DNA was precipitated from the supernatant with isopropanol and washed 3 times with 70% ethanol. Purified DNA was stored at -20 °C for future use.

**Agilent SureSelect hybridization enrichment and sequencing**

Nuclei were isolated from human prefrontal cortex via FANS. Genomic DNA was extracted and fragmented using sonication (Covaris, Woburn, MA). End repair reactions were performed and Illumina sequencing adaptors were ligated to genomic DNA. Library-prepped DNA was hybridized with Agilent SureSelect probes designed against all human exons (V6 without UTRs). Pulled-down genomic DNA sequences were then sequenced on an Illumina NextSeq (Illumina, San Diego, CA).

**Identification of *APP* gencDNAs and insertion sites from 10 pull-down datasets**

Fastq files from the Park *et al*.^2^ datasets were downloaded from SRA (accession PRJNA532465). Sequences from the Park *et al*. and Chun Lab datasets were first aligned to the human reference genome (GRCh38) using BWA-mem (version 0.7.17) and screened with Vecuum (version 1.0.1) to confirm that APP plasmid was not detected in any of these datasets. Datasets were then aligned to the human genome using STAR (version 2.7.3a) with the options previously described in Lee *et al.^3^* (--outSAMattributes All --outSJfilterCountTotalMin 1 1 1 1). We excluded reads that contained the tag ‘jI:Bi:i,1’ using SAMtools (version 1.3) in order to keep the reads where a junction was detected. To remove reads that were “mis-split” by STAR, we filtered out reads that had fewer than 16bp mapped to each side of the junction (at least 16bp of each exon are required to improve mapping accuracy). Duplicate reads were marked and removed using Picard (version 2.1.1). Reads were then processed and visualized using a modified version of the R exonjunction package (<https://github.com/christine-liu/exonjunction>).

To identify APP novel insertion sites, we first extracted discordant read pairs in which one read mapped to an APP exon::exon junction or was clipped (>= 20 bps) at APP UTR ends, and the mate read mapped confidently (mismatches < 1%) to a novel chromosome other than chromosome 21. We also made sure that these reads mapped uniquely to the human genome but not to the mouse genome or transcriptome. Next, we mapped the UTR clipped sequences against the human genome using BLAST+ (version 2.9.0) and kept only the reads confidently aligned (query coverage >= 90%, identical matches >= 98%) to the same chromosomes as their corresponding mates. Detailed read alignment information can be found in Supplementary Table 1.

**Supplementary Discussion of gencDNAs in blood**

The source of blood gencDNAs is not known but could be from lymphocytes that can express all *APP* forms*^4^*, or could suggest that neural gencDNAs can be detected in blood and may have value as a potential biomarker.

**Supplementary Discussion of DISH and RISH**

DISH and RISH are important to our conclusions and cannot be ignored. Our DISH and RISH probes were custom designed with Advanced Cell Diagnostics (ACD, Newark, California) to target the exon::exon junction between exon 16 and 17 and the IEJ between exon 3 and 16. Multiple controls and experiments validated the specificity of our ISH probes (*Lee* Figure 2j-n, Extended Data Figure 1f, 2f-h, 6b, 6d-f)^3^: 1) sense and antisense probes that demonstrated DNA specificity of the sense-strand probe; 2) targeted restriction enzyme digestion that effectively destroyed the target locus and eliminated probe-dependent signal; 3) double-labeling that showed gencDNAs were usually distinct from the endogenous locus; 4) creation and use of retrovirally-delivered synthetic targets in cell culture that confirmed probe specificity; and 5) use of the J20 mouse model that contains the human *APP* sequence that confirmed specificity by the sense-strand identification of the transgene. ACD and others have demonstrated the ability of BaseScope^TM^ probes to differentiate exon::exon junctions^5^, splice variants, and point mutations^6^, and therefore, we are confident that our RISH and DISH probes identify *APP* gencDNAs and IEJs in a PCR-free and plasmid-free system. Additionally, DISH resulted in cell type and disease-specific patterns of gencDNA expression, with SAD neurons showing the greatest number of gencDNA foci, while J20 mouse experiments demonstrated specificity as shown in Lee *et al.* None of these results could be explained by PCR of *APP* plasmid sequences.
